# Biophysical basis of alpha rhythm disruption in Alzheimer’s disease

**DOI:** 10.1101/335471

**Authors:** Rohan Sharma, Suhita Nadkarni

## Abstract

Alpha is one of the most prominent rhythms (7.5–12.5 Hz) detected in electroencephalography (EEG) during wakeful relaxation with closed eyes. In response to elevated ambient acetylcholine levels, a subclass of thalamic pacemaker cells generate alpha. This rhythm is intrinsic to the cell and is robustly orchestrated by an interplay of hyperpolarization activated cyclic nucleotide gated channels(HCN) and calcium-ion channels. It has been shown that decreased expression of HCN channels is correlated to Alzheimer's Diseased (AD). In early stages of AD, alpha is known to be down-regulated and lowered in coherence. We use this well characterized and quantified rhythm to understand the changes in ion channel properties that lead to disruption of alpha as seen in AD in a biophysically detailed network model of the thalamo-cortical circuit that generates the alpha-rhythm. Our computational model allows us to explore the causal links between alpha rhythms, HCN channels and amyloid-beta aggregation. The most commonly used drugs(acetylcholinesterase inhibitors) in AD increase the duration and level of acetylcholine and provide temporary symptomatic relief in some cases. Our simulations show how increasing acetylcholine can provide rescue for a small range of aberrant HCN expression. We hypothesize that reduced alpha rhythm frequency and coherence is a result of down-regulated HCN expression, rather then compromised cholinergic modulation(as is currently thought). The model predicts that lowering of the alpha-rhythm can modify the network activity in the thalamo-cortical circuit and lead to an increase in the inhibitory drive to the thalamus.

## 1 Introduction

It is widely agreed that rhythms play a vital role in coordinating and organizing neuronal computations across various anatomical regions of the brain. Robust and sustained rhythms, from a fraction of a hertz (delta) to several hundred hertz, have been implicated in a range of functions, like attention, spatial navigation and memory consolidation. The alpha-rhythm in particular has been associated with the cognitive function of attention (selection and suppression) and semantic orientation [1]. Typically, network mechanisms that invoke interplay between inhibitory and excitatory cells lead to genesis of rhythms that are robust as well flexible in response to ongoing functional requirements [2]. Gamma rhythms, for instance, are generated by synaptic interactions between inhibitory neurons and excitatory pyramidal neurons, the so called pyramidal interneuron gamma (PING) [3]. Alpha rhythms, on the other hand, are an intrinsic property of neurons in the thalamus; blocking chemical synaptic transmission does not block alpha [4]. The alpha rhythm generated in the thalamus is the result of synchronous bursting of neurons, each firing at a frequency within the alpha range. They maintain stable phase relationships because of tight gap junction coupling. Studies also suggest that cortical neurons too can generate alpha [5]. Alpha-rhythm activity has also been observed in other brain regions like the the pre-frontal [6], auditory [7] and somatosensory cortex [8]. In response to a discrimination task a reduction in the amplitude of the peak at 10 Hz is observed, along with the emergence of a low amplitude peak at 20 Hz [9]. This suggests that the coherence and frequency changes in alpha are functionally relevant attributes.

Here we have used a biophysically detailed, conductance based network model of neurons in the thalamus associated with alpha generation [10]. The model comprises of a set of specialized thalamic cells, with a high threshold calcium current (HTC cells), which fire at alpha frequency(10 Hz) during high levels of ambient acetylcholine that activate muscarinic acetylcholine receptor(mAchR) [4]. This is modeled as a lowered potassium leak conductance *g_Kleak_* Eq:1 [10, 11]. Each of these cells are connected via gap junction to ensure synchronous activity. Even though the rhythm generation is limited to the HTC cells, they are not isolated from the rest of the thalamic network. Our model utilizes a minimal network motif which follows physiologically realistic ratios of inputs and outputs associated with the thalamus 1, including the HTC cells and their synaptic interactions. This minimal network allows us to explore the effects of the HTC cell firing on the rest of the network and the effect of the network firing on the HTC cells. It consists of two HTC cells connected to each other via a gap junction, which provide an excitatory drive to the RE cells and TC cells. The TC cells are excitatory and connect to all RE cells but not to each other or the HTC cells. The RE cells are GABAergic and inhibit every other cell in the network, including other RE cells 1.

**Fig 1.**
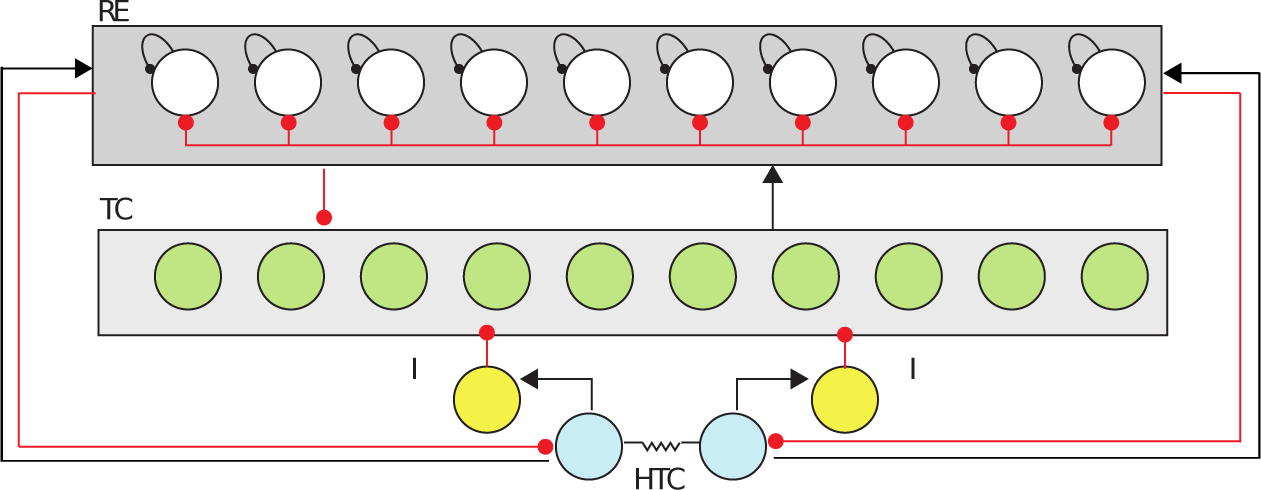
Thalamocortical Network involved in Alpha generation. The RE cells (white) mutually inhibit each other and inhibit all of the TC cells (green). The TC cells send excitatory connections to all of the RE cells. There are no direct mutual connections between TC cells. The HTC cells (blue) are solely responsible for generating the alpha rhythm inhibit the TC cells via sending excitatory drive to the inhibitory-interneurons (yellow). The ratio of the RE to TC cells is 1:1 and 20% of the TC cells are HTC cells. The inhibitory interneurons are modeled implicitly, using a synaptic delay. Gap-junctions between the HTC cells enable synchronized bursting.

The causal link of events that chronicle the activity of ionic currents pivotal to the generation of alpha-activity, leading to initiation and termination of the rhythm in the presence of acetylcholine is described in 2. High threshold calcium ion channels and a nonspecific (for ions) hyperpolarization activated cyclic-nucleotide gated(HCN) channels are the ionic components of HTC cells that govern alpha. The precision and robustness of the rhythm is determined by the intrinsic dynamics of the gating variable of each current and the interplay between the currents mediated by membrane voltage and calcium. After each burst of action potentials, the cell undergoes a brief hyperpolarization, which activates the HCN channels. The slow positive inward current due to opening of HCNs then steadily depolarizes the cell till the opening threshold of the calcium channel is reached. This calcium channel is favored with an instantaneous activating rate and a slow inactivating rate. They also have a narrow regime of voltage ~(−50 mV to −10 mV) over which they are open. The calcium current causes rapid depolarization, triggering the fast sodium and potassium channels in the presence of high levels of acetylcholine (*g_Kleak_*) and generate a series of action potentials. The sluggish response of inactivating gate of the calcium channel to the ongoing action potential causes a slow decay of this current. The decay eventually terminates the action potential burst, allowing potassium channel to repolarize the membrane. As the membrane repolarizes, the HCN channels start getting activated, setting up the system for another burst. The depolarization time scale sets the time interval between bursts and defines the alpha rhythm. The depolarization time scale is determined by the magnitude of the I_H_ current, which in turn depends on the HCN channels’ conductance and expression.

Neurological disorders like Alzheimer’s Disease (AD), other forms of dementia, Parkinson’s disease etc. all have have characteristic signatures in EEG recordings [12]. AD patients in particular show diminished power and down-regulated frequency in the alpha band [13] [14]. In the computational model described here and in slice studies [4], the alpha frequency and power is keenly modulated by ambient concentration levels of acetylcholine. A class of drugs that inhibit the breakdown of acetylcholine(acetylcholinesterase inhibitors) and therefore augment its resting levels, provide temporary symptomatic relief. These observations portend an interesting correspondence of alpha, its disruption in AD and its downstream cognitive implications.

Individuals with a genetic risk of AD (APOE-4 carriers) have been shown to exhibit reduced grid-cell-like representation and have difficulties in navigating in familiar environments [15]. Grid cell representation in the entorhinal cortex has been shown to have a gradient along the dorso-ventral axis. As we move along the axis, the grid cell spacing increases. The grid cells themselves show a gradient in the bio-physical properties of the HCN1 channels. The HCN1 channels’ expression and the time constant of activation increases along the dorso-ventral axis, while the size and spacing of grid cells increases in HCN1 knockout mice [16]. This association between grid cells and the bio-physical properties of HCN1 channels, suggests a close association of HCN1 channels with grid cells. We believe disrupted expression of HCN1 channels can be detrimental to the cognitive ability of a subject to perform spatial navigation. This makes us believe that the initial loss in the ability to perform spatial navigation in early stages OF AD arises out aberrations in the expression of HCN1 channels in the entorihnal cortex. Indeed lowered activity of HCN1 channels has been shown to increase the production of A*β* from amyloid precursor proteins(APP) [17]. This suggests decreased amount of HCN channels could be an upstream event to A*β* production in AD neurons. There might lie an explanation of observed mild cognitive decline even before more prominent markers associated with AD like A*β* and Neurofibrillary Tangles(NFT) are seen [18]. We systematically explore the consequence of reduced HCN channel activity on the alpha rhythm. Prompted by the therapeutic merit of acetylcholinesterase inhibitors and the role of acetylcholine in initiating the alpha rhythm, we also investigate the rescue of the alpha rhythm by increasing cholinergic modulation.

**Fig 2.**
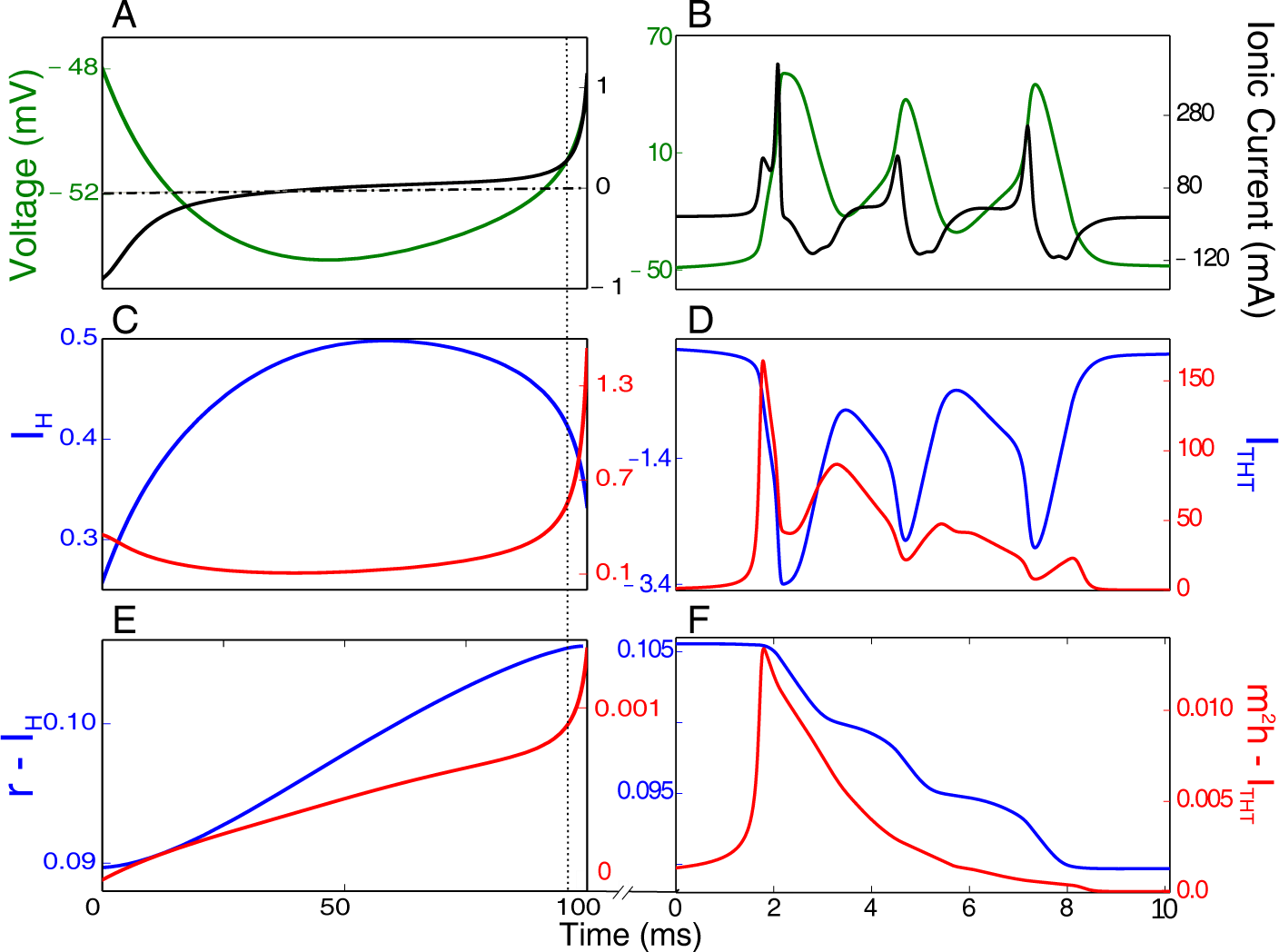
Chronology of events underlying the alpha-rhythm. The bursting activity in alpha-rhythm is initiated by activation of *I_H_* current (B) and terminated by repolarization with the potassium channel. The inter-burst interval determines the 10 Hz rhythm. The left panels describe slow dynamics between the burst over a 100 ms time window. The right panels describe activity of ion channels during a burst over 10 ms. (A) Slowly increasing membrane voltage (green) due to increase in *I_H_* conductance (during repolarization of the membrane by potassium channel). The total current (black) increases with *I_H_* B) Time trace of the *I_H_* current (blue) as it slowly activates during repolarization leading to the activation of the *I_THT_* (red) close to the 90 ms mark(dotted line) C) Gating variables of the *I_H_* (*r_H_*) and *I_THT_* (*m*^2^*h*) currents. The high-threshold calcium current gets activated around the 90 ms mark(dotted line) D) Zoomed in to the the first 10 ms after of activation *I_THT_*. Burst of action potentials are generated driven by membrane depolarization initiated by *I_H_* and *I_THT_* in that order. E) The high-threshold calcium current provides the depolarizing impetus that sustains a burst of action potentials. F) The slow decay of the gating variables of the *I_H_* as the membrane potential rises above the activation voltage. The gating variables of *I_THT_* slowly deactivate at the high voltages during action potentials.

## Materials and Methods

### Details of ionic currents associated with the neuron, synapses and the network connections in the thalamo-cortical circuit that generates the alpha-rhythm

The model consists of the canonical point-neuron network model of the thalamo-cortical circuitry with the addition of a high-threshold T-type calcium current in 20% of the TC cells (HTC)1. The HTC cells receive a white-noise input with zero mean which was implemented using the Euler–Maruyama method. The TC and RE neurons receive a poisson-distributed train of excitatory and inhibitory impulses. The activation of the mAch receptors is phenomenologically modeled by lowering the potassium leak conductance [19] We used an in-house Computational Neuroscience library written in C++ https://github.com/insilico-lib/insilico to do the simulations. The time step of each simulation was taken to be 0.01 ms.

### Neurons

#### Thalamo-reticular(RE) neurons

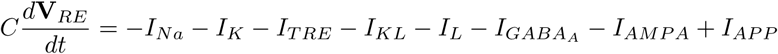

Potassium Current:

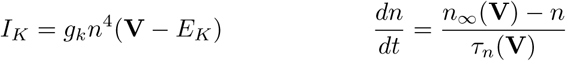

here:

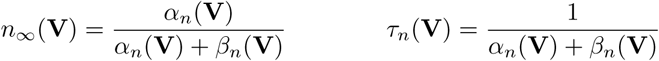

where:

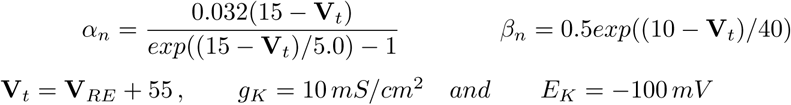

Sodium Current:

The *m_∞_, τ_*m*_, *h*_∞_ and τ_*h*_* have equations identical to the *n_∞_* and the *τ_n_* of the potassium gate n.

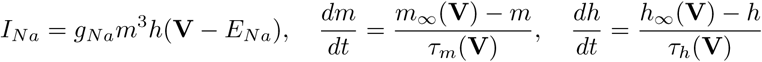

where:

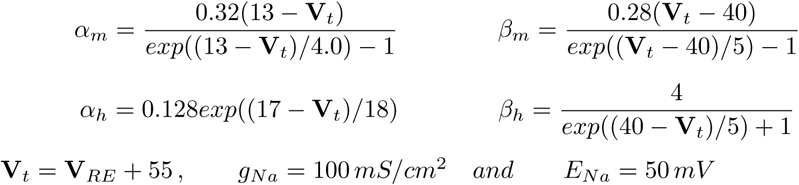

Calcium Current:

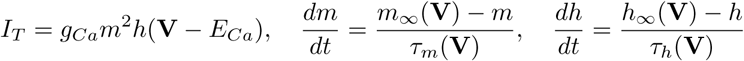

where:

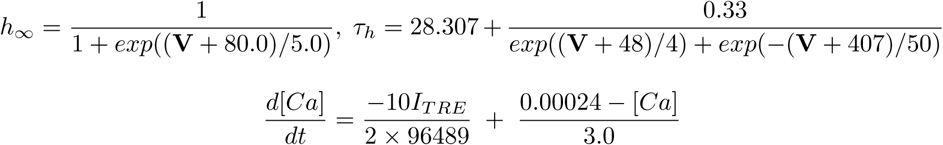

The first term must be positive, otherwise it is set to zero. *g_Ca_* = 2.3*mS/cm*^2^ and the reversal potential for calcium is calculated using the Nernst-Equation

Leak Current:

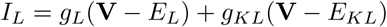

where:

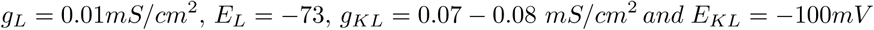

Applied Current:

The applied current is a train of poisson-distributed excitatory and inhibitory impulses. The details of the same will be discussed later.

#### Thalamo-cortical(TC) neurons

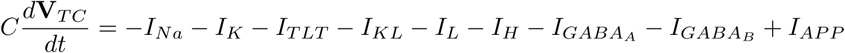

Potassium Current:

It is very similar to the RE cell potassium current except:

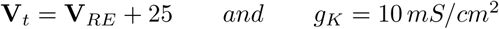

Sodium Current:

It is very similar to the RE cell sodium current except:

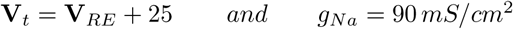

Low Threshold Calcium Current:

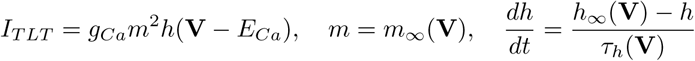

where:

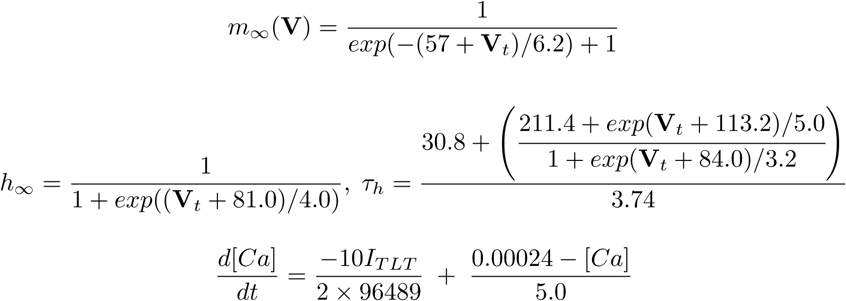

The first term must be positive, otherwise it is set to zero.

*g_Ca_* = 2*mS/cm*^2^, **V***_t_* = **V** + 2 and the reversal potential for calcium is calculated using the Nernst Equation

Leak Current:

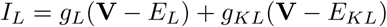

where:

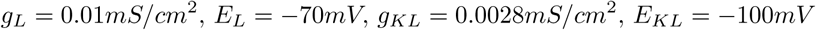

H-Current:

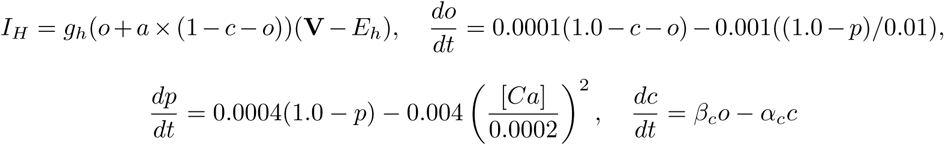

where:

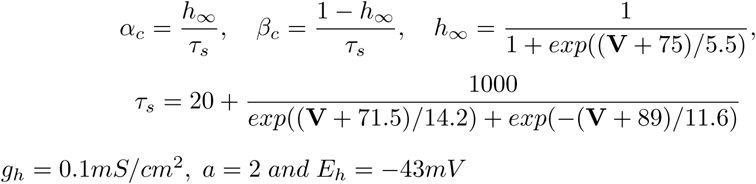

Applied Current:

The applied current is a train of poisson-distributed excitatory and inhibitory impulses. The details of the same will be discussed later.

#### High Threshold Thalamo-cortical(HTC) neurons

Potassium Current:

The potassium current follows the same dynamics as the potassium current in TC cells

Sodium Current:

The sodium current follows the same dynamics as the potassium current in TC cells

Low Threshold Calcium Current: The low threshold calcium current follows the same dynamics as the potassium current in TC cells

High Threshold Calcium Current:

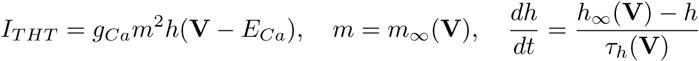

where:

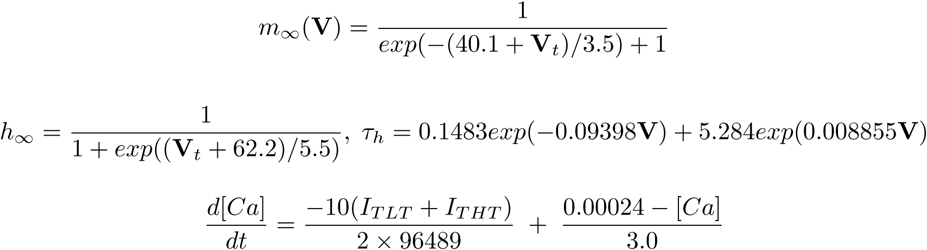

The first term must be positive, otherwise it is set to zero.

*g_Ca_* = 12*mS/cm*^2^ and *E_Ca_* is calculated using the Nerst-equation

Leak Current:

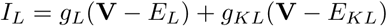

where:

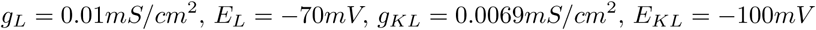

H-Current:

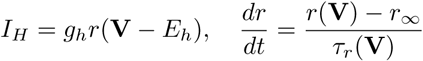

where:

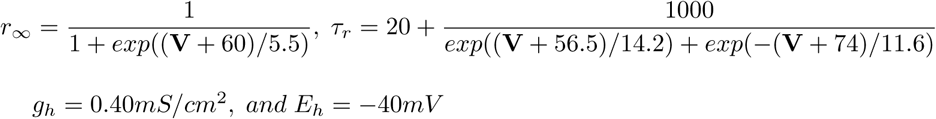

Calcium Activated Potassium Current:

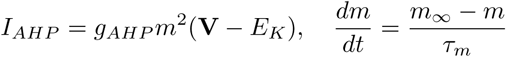

where:

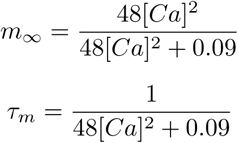

*g_AHP_* = 15*mS/cm*^2^ and *E_K_* = *−*100*mV*

Gap Junction Current:

*I_GJ_* = *g_GJ_* (**V***_HTC_* − **V***_post_*), where **V***_post_* is the membrane potential of the neuron that is connected to this HTC neuron by a gap junction

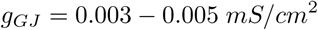

Applied Current:

The HTC neurons receive a Gaussian white noise with a mean around zero and standard deviation of 0.1

#### Synapses

AMPA:

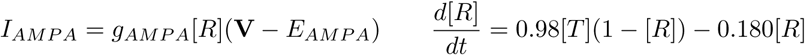

[T] is the transmitter concentration. When a pre-synaptic cell experiences and action potential the transmitter concentration is increased to 0.5 mM and stays there for 0.3 ms for HTC and 0.5 ms for TC cells. [R] represents the fraction of the receptors that are open.

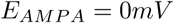

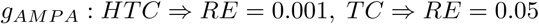

*GABA_A_*:

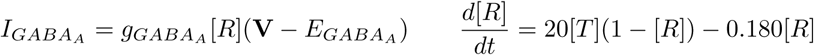

[T] is the neuro-transmitter concentration. When a pre-synaptic cell sees an action potential the transmitter concentration is increased to 0.5 mM and stays there for 1.0 ms for HTC and 0.3 ms for RE cells. [R] represents the fraction of the receptors that are open.

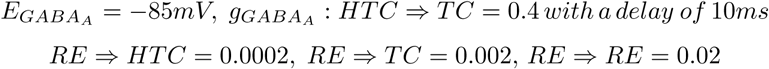

*GABA_B_*:

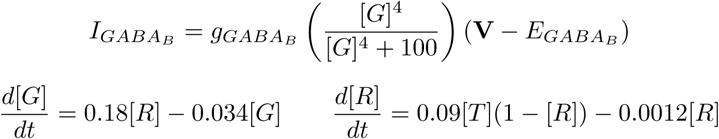

[T] is the neuro-transmitter concentration. When a pre-synaptic cell sees an action potential the transmitter concentration is increased to 0.5 mM and stays there for 0.3 ms. [R] represents the fraction of the receptors that are open. [G] is the concentration of the G protein that gets activated upon agonization of the receptors.

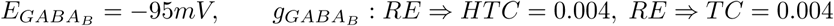

*Noise*:

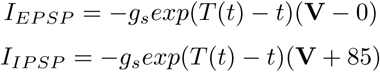

where:

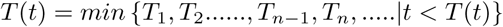

The difference between the impulse times, *T*_1_, *T*_2_.....*T_n_* is an exponentially distributed random variable with a mean of 10 ms for RE cells, which have *g_s_* = 0.02*mS/cm*^2^ for EPSPs and *g_s_* = 0.015*mS/cm*^2^ for IPSPs. For TC neurons the mean is also 10 ms for EPSPs with *g_s_* = 1.0*mS/cm*^2^, but they are not given any IPSPs.

The HTC cells receive a gaussian distributed white noise through the stochastic Euler-Maruyama integrator:

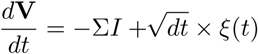

where *ξ*(*t*) is drawn from a Gaussian distribution with mean 0 and variance 0.1

##### Entropy Measure

V_i_ is the membrane voltage of the HTC neurons, where *i* ∈ {1, 2}

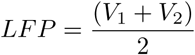

Before performing the Fourier transformation, we do some basic processing over the LFP trace. We take a simple moving average over a window of 10 ms(we use an observation frequency of 2.5kHz) and make it’s mean zero.

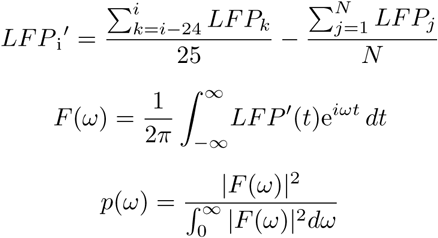

*p*(*ω*) is the the probability distribution function which is used to calculate the Shanon Entropy

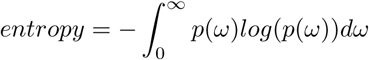

## Results

### Effect of varying HCN expression on alpha

Diminished expression of HCN channels has been reported in Alzheimer’s effected neurons [17]. It has also been shown that lowered HCN expression can accelerate amyloid-beta aggregation. HCN channels are crucial in initiating the alpha rhythm (see Fig2 2). To investigate the disease pathology, we explore the effect of aberrational HCN expression on the alpha-rhythm by modulating the conductance of the I_H_ current(g_H_). Control value for the conductance set to around ~0.36mS/cm^2^ in the model gives robust periodic alpha at 10 Hz [10]. We quantify the periodicity using the entropy of the FFT of the local field potential(LFP). Probability distribution function (pdf) for the entropy measure is represented by normalized power spectrum of the LFP(see methods section for details). Higher values of the entropy imply that the power in the signal is distributed over various frequencies. A neatly periodic and coherent time series will exhibit low entropy, as the power would be confined to only a few narrow frequency regions. For example, see 4CI. (Supplementary information S1 Fig) shows the relationship between this entropy and variance of peak frequency.

Increasing g_H_ increases the frequency of HTC firing monotonically(−25% to + 20% change in g_H_) (3A peak frequency in power spectra in blue). Beyond this regime of g_H_, periodicity breaks down and alpha rhythm is lost. This is seen as increase in entropy in both directions of control value of g_H_. Increased expression of I_H_ (higher value of g_H_) makes the membrane more excitable so that small fluctuations are more likely to cross threshold causing noisy firing. Decreased expression of I_H_ (g_H_) launches I_THT_ later. These changes in g_H_ have the effect of reducing the intrinsic bursting timescale of HTC cells as described in the introduction and moving out of the alpha range. As a result the HTC activity rate is seen to slow down until g_H_~0.27mS/cm^2^. At values of g_H_<0.27mS/cm^2^, HTC cell membrane is just below firing threshold over longer periods of time making the system again sensitive to noise. Background noise has longer periods of opportunity to cause the crossing of firing threshold. Our calculations suggest a high sensitivity of the alpha rhythm to the HCN expression and a narrow regime of HCN expression over which periodic activity of HTC cells is possible.

**Fig 3.**
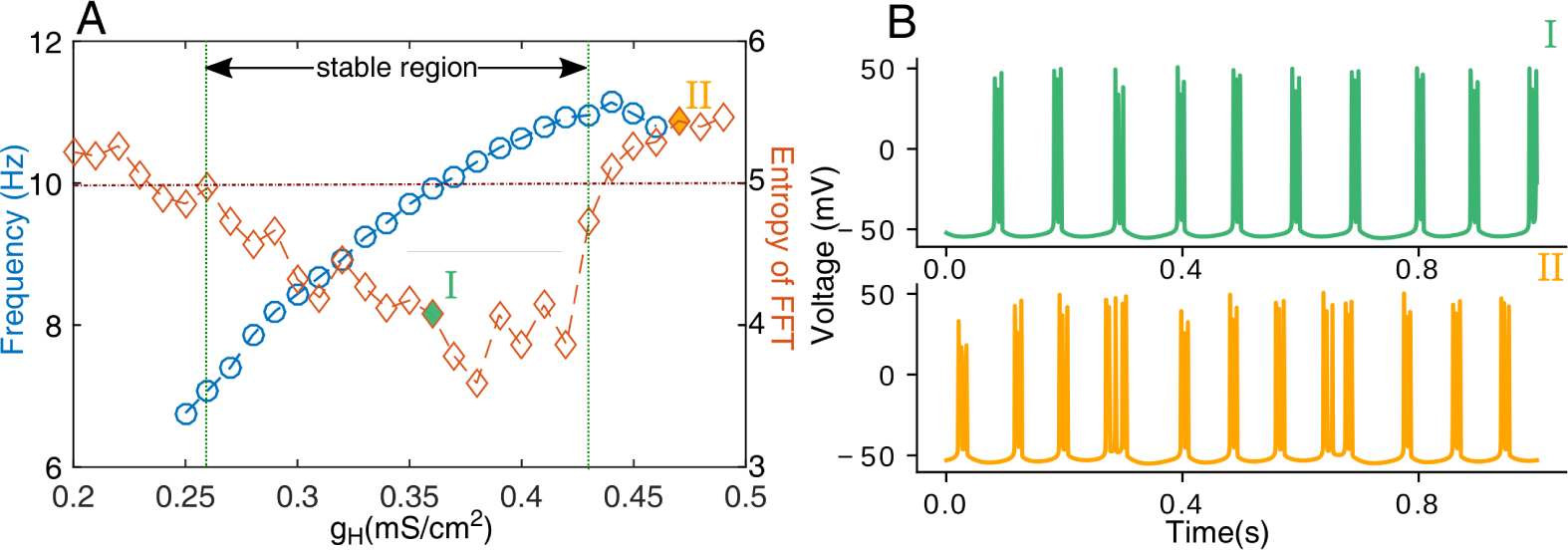
Monotonic dependence of the peak frequency by varying HCN expression. There is small range of g_H_ values that will give rise to a periodic bursting phenomenon. **(A)** Intrinsic oscillation frequency (blue) of HTC cells show monotonic dependence on conductance g_H_ of HCN channel in the stable region. However I_H_ over expression leads to over excitability in HTC cells and losing periodicity. Low levels of I_H_ expression makes the system more sensitive to noise and loss of periodicity. There exists an optimal regime of g_H_ where bursting is regular. This is depicted as lower entropy (red) for a range of g_H_. **(B)** Illustrative voltage traces of HTC cells from I(g_H_=0.36mS/cm^2^) and II(g_H_=0.47mS/cm^2^)

### Limited rescue of the alpha-rhythm with acetylcholine

Ambient acetylcholine acts to increase global excitability of HTC cells. This trigger, along with the intrinsic properties of these cells, initiates alpha. In AD, reduced HCN expression in HTCs leads to lower excitability and delays the activation of the burst inducing calcium-current. The chain of events that determine alpha-rhythm time-scale now take a longer time to complete. Here we investigate the extent to which increased acetylcholine levels can counter the lowered excitability and reinstate alpha. In figure 4 four case studies of differential HCN expressions (Green: normal, Blue: Increased expression, Black: reduced) and its implications on alpha rhythm are described. We define *η_ach_* as a measure of fractional change in cholinergic activity.

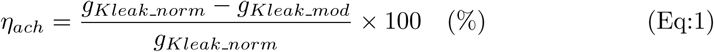

Where *g_Kleak_norm_* is the potassium leak which corresponds to a 10 Hz alpha rhythm and *g_Kleak_mod_* is the modified potassium leak. Increase in ambient acetylcholine levels (*η_ach_*)lead to monotonic increase in alpha frequency over a limited range. As the system of HTC cells continue to get more excitable with *η_ach_* they inadvertently also become to susceptible to noise. Normal HCN expression maintains periodicity upto around 15% increase in *η_ach_* (green diamonds). HTC cells with reduced HCN expression of ∼80% (black diamonds, fig 4A) can tolerate increase up to 28% increase in *η_ach_*, beyond which there is a loss in periodicity. This is illustrated in Fig 4B bottom figure. The transition for cells with lower HCN expression to normal alpha rhythm (rescue) happens at an increased value of *η_ach_* = 20%. The loss in periodicity is quantified as an increase in entropy, as defined earlier (filled diamonds). Changes in power distribution around the alpha band with the healthy, pathological and rescue cases of the alpha-rhythm are shown in 4. Our results suggest that rescue by acetylcholine is possible in a limited range of reduced HCN levels. Further increase in excitability, with higher value *η_ach_* lead to enhanced sensitivity to noise. Figure 4C describes HTC rhythm and corresponding entropy for HCN levels reduced to 45%. For HCN levels as low as these, we see that increases in acetylcholine cannot bring the HTC rhythm up to 10 Hz and periodicity is restricted t0 9 Hz, beyond which the HTCs lose rhythmicity (see 4C).

**Fig 4.**
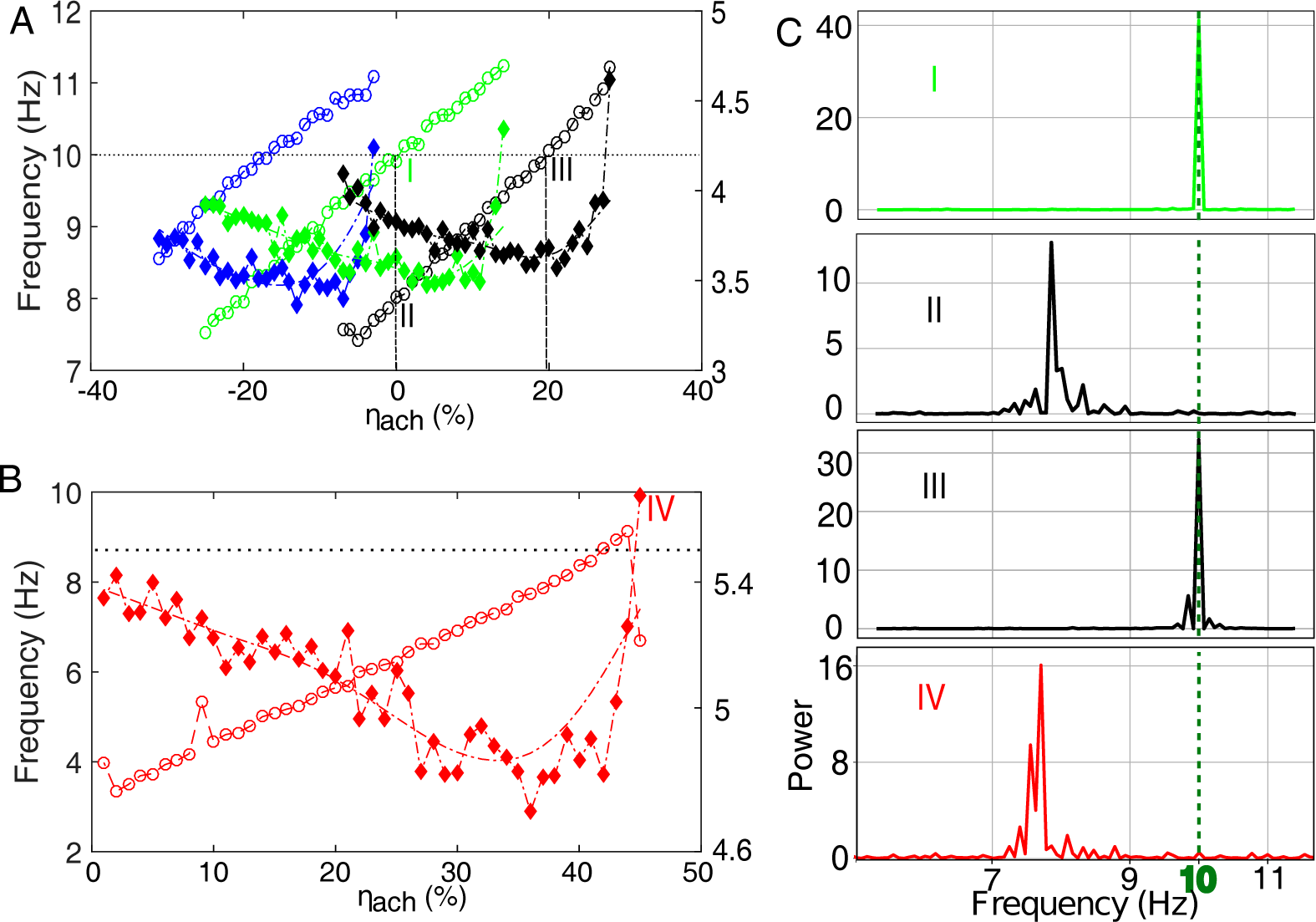
Rescue of the alpha-rhythm to changing levels of basal [Ach] and HCN channel expression. (A) Increasing ambient acetylcholine levels (*η_ach_*) increases the peak frequency of the rhythm (open circles, blue, green and black). When I_H_ expression is compromised (78% black open circles) to mimic known observations in AD, peak frequencies remain lower than control (100% g_H_, open green) (II). The lowered coherence of alpha as a result of lower g_H_ can be rescued by increasing acetylcholine levels (78% g_H_ expression needs 20% increase in acetylcholine(III). There appears to be a threshold of ambient acetylcholine levels beyond which entropy(filled diamonds) increases dramatically, suggesting loss in periodicity. The overall entropy also remains high for decreased g_H_. (B)When I_H_ expression is compromised severely (50% green open circles) lowered frequency of alpha is not rescued by increasing acetylcholine levels. 50% g_H_ expression can only achieve periodicity 9 Hz before complete breakdown of regular firing. This is seen as a sudden rise in entropy (closed red diamonds). (C)Power Spectra corresponding to I, II and III from (A) and IV from (B).

### Lowering g_H_ can lead to an enhanced GABA activity

Several studies have characterized compromised alpha in AD [20]. So far we have shown how changes in acetylcholine levels and HCN channel can modulate alpha. Here we investigate how AD related changes in alpha can influence network dynamics. During alpha band activity, rest of the brain areas that are not corralled into the oscillations, are suppressed [21]. This leads to the notion that alpha activity inhibits neuronal firing. Using the computational model described above we analyze the effect of changing the alpha frequency on the dynamics of the thalamocortical network. To mimic the AD condition we diminish HCN expression. As expected, the frequency of HTC firing is seen to decrease with lowered I_H_ conductance(g_H_) 3A. Under these pathological conditions with lowered alpha, the TC cells that were suppressed via GABAergic drive from the rhythm generating HTCs are now released from inhibition 1&5. In figure 5A we show increased TC firing as a result of lowered HTC frequency. The reduced inhibition from HTCs seems to have a downstream effect on RE cells. The RE cells which receive enhanced excitatory drive from TC cells, transition to higher firing rates. The increased response of RE cells to increased excitatory drive from the TC cells is shown in 5B. This happens despite the decreased drive from HTCs to the REs. HTC cells are 20% of the total number TC cells(normal TC cells and HTC cells). The larger number of TC(4 times) overrules the synaptic interaction and creates a positive feedback loop. High RE activity leads to increased inhibition on the HTC cells reducing their frequency further 5. The overall effect is that of increase in GABAergic activity. In support of this insight, increased presence of neurotransmitter GABA has been reported in AD mice [22]. Our model suggests that lowered alpha rhythm seen in AD can cause a cascade of changes in thalamocortical network firing and may ultimately cause increased inhibition.

**Fig 5.**
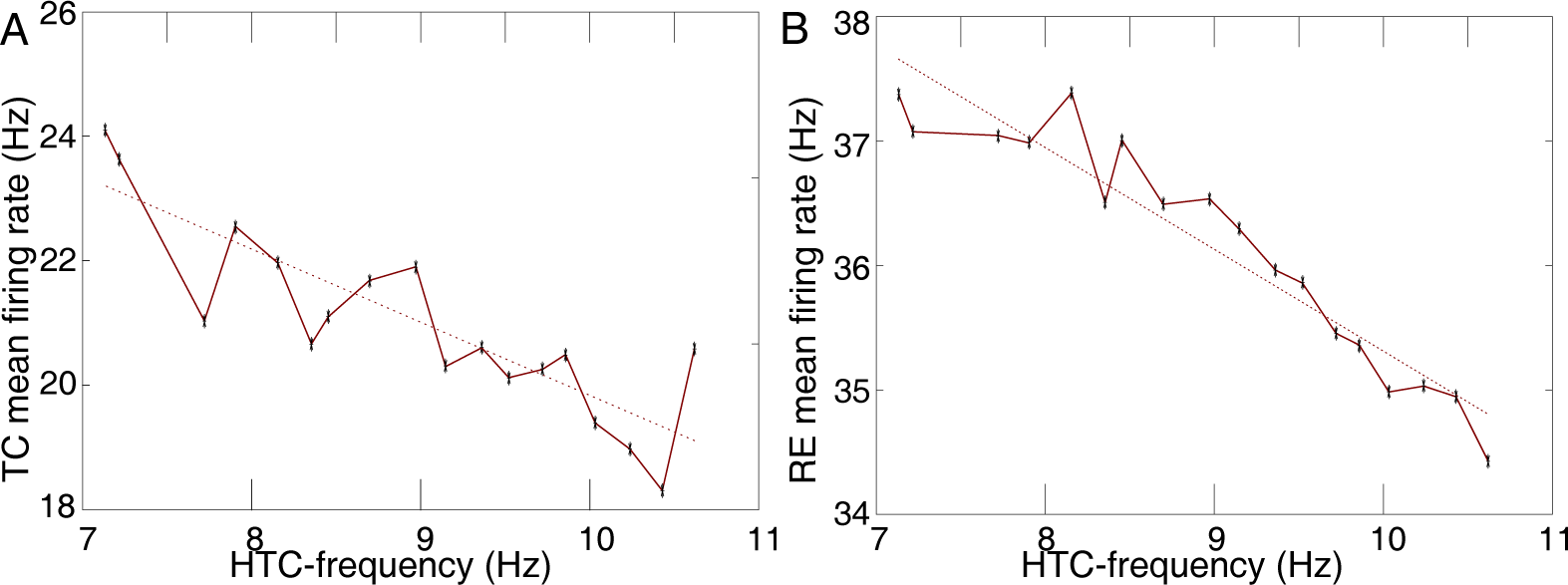
Increase in inhibitory activity follows a reduction in the HTC-frequency. A) Inverse relationship between HTC and TC firing. Lower drive from HTCs releases the inhibitory drive (via interneurons) on the TC cells causing increase in activity. B) Inverse relationship between HTC firing and RE firing. Increased activity of the TC cells leads to enhanced RE activity.

### Calcium current and its interaction with HCN expression

Disrupted calcium homeostasis is been implicated in several studies in Alzheimer’s affected neurons. Elevated levels of cytosolic calcium are associated with AD and linked to dysregulation in the calcium signaling within the cell [23]. We simulate the pathology by changing the high-threshold calcium conductance in the cell (crucial for alpha). Interestingly, despite the crucial role of calcium in orchestrating the rhythm, individual HTC cells fire in a small range around alpha in response to as much 50% to 150% of normal high-threshold calcium conductance (g_THT_). This surprising readout is due to the narrow frequency repertoire of the HTC cells. However our model suggests that disruption in alpha caused by changes in calcium arises out of loss in periodicity. In other words, HTC cells either fire around alpha or loose periodicity. In order to isolate the effect of the calcium conductance on the alpha-rhythm, we change potassium leak to strictly maintain the 10 Hz frequency for the range of *g_H_* explored in the figure. As calcium conductance is changed, HTC cells go through regimes of periodic and irregular firing. This is shown as sudden transitions in entropy in 6A. Healthy cells (g_H_=0.36mS/cm^2^, 6A, green) appear robust and can tolerate as much as a 25% increase in the calcium conductance before losing periodicity. On the other hand decreasing g_H_ values to simulate the pathological condition of lowered HCN expression shows lower tolerance for changes in calcium conductance. This heightened sensitivity to calcium is shown as narrower troughs in entropy (windows of regular firing) for pathologically lower expression of HCN shown in orange and blue. 6A also describes response to increase HCN (red and purple). The overall effect follows the same trend of increased tolerance to changes in calcium which corresponds with HCN expression. We summarize this in 6B. The entire range from −50% to + 50% change in calcium conductance is shown. Lowered HCN expression corresponds with shrinking regimes of periodic firing 6.

**Fig 6.**
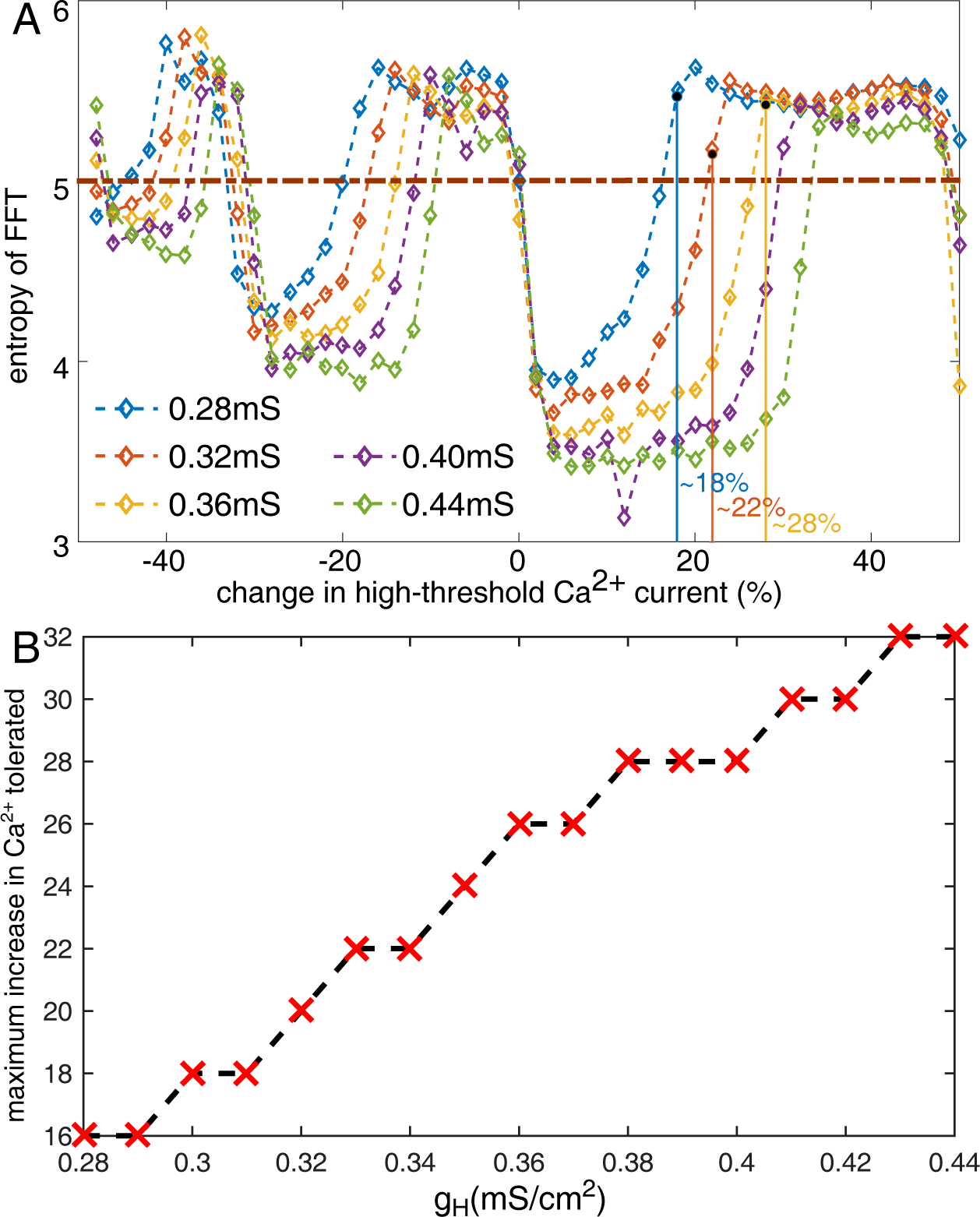
Lowered HCN makes Alpha more sensitive to small changes in the calcium conductance. A) Systematically varying the calcium conductance in both directions leads to sudden increases in the entropy describing incoherent firing of the HTC cells. These are regimes of calcium conductance over which the HTC cells show aperiodic activity. Coloured lines show deviations from normal HCN expression levels. Lower HCN expression has lowered tolerance for changes calcium. Higher HCN gives a broader range of calcium conductance where we see periodic firing. B) Finer illustration of higher sensitivity of pathological HCN expression to changes in calcium.

## Discussion

Alzheimer’s disease(AD) is a multifaceted catastrophic disease that implicates multiple brain areas, resulting in a range of debilitating symptoms, making it difficult to arrive at a nodal cause. While precise molecular mechanisms that underlie the constellation of deficits and the causal links between them are not completely clear, however biochemical and electrophysiological markers have been observed. Several apparently independent hypotheses have been proposed to delineate the root pathology and the consequent pathogenesis.

The most prominent of these is toxic effects of accumulation amyloid-beta plaques and Tau fibrils, a characteristic feature of AD [24]. While amyloid-beta and tau-fibrils disrupt a wide array of signaling pathways in the brain, that include cell death, we do not yet have a clear understanding of the biochemistry that leads to their accumulation and proliferation. It has been suggested that reduced HCN expression and down-regulated HCN channel activity could be leading to increased amyloid-*β* aggregation.

The calcium hypothesis of Alzheimer’s suggests that dysfunctional regulation of the calcium signaling by modifying synaptic plasticity and other signaling cascades, profoundly vitiates neural functions like memory formation and consolidation. However its not known how small changes in the calcium signaling cause drastic changes in behavior [23]. We demonstrate in this work how changes in the parameters that govern calcium dynamics can lead to irregular activity in the thalamus 6. Reduced HCN expression, associated with AD, can make the network more sensitive to deviations in calcium signaling.

The cholinergic hypothesis proposes reduced release acetylcholine as the leading cause of symptoms of AD. In support of this, the most prevalent drugs administered to AD patients that provide temporary symptomatic relief are acetylcholinesterase(Ache) inhibitors [25, 26]. EEG tools that are used to diagnose AD, report a reduction in power and frequency of alpha compared to control subjects [27] [13]. This evidence suggests compromised acetylcholine signaling in AD. Results described here quantify the differential ways by which changes in ambient acetylcholine can modulate this rhythm. However in light of the role of HCN expression in AD and the insights from the model, we hypothesize that it is not aberrant acetylcholine signaling itself that is the cause of AD symptoms.

Alpha is essentially orchestrated by action of ambient acetylcholine that depolarizes the membrane and in turn gets HCN channels and calcium channels to generate the characteristic 10 Hz burst. Reduced HCN expression and over-expression of beta amyloid are key observations in AD cells [17]. We clearly demonstrate how reduction of HCN channel expression can make the cell more susceptible to background noise with minor deviations in the calcium current 6. In view of reduced power and coherence seen in the alpha band of AD patients, our predictions connect changes in calcium to aberrant HCN expression in AD. Using some of the known observations linked to AD and a biophysically detailed computational model that generates alpha, we illuminate the possible causal relationships between key markers associated with AD (Beta Amyloid, HCN expression and Acetylcholine hypothesis) 7.

### Analysis of causal relationships between HCN expression, alpha-rhythm and amyloid-beta aggregation

In AD transgenic mice cognitive decline is observed before amyloid-beta plaques are visible. Drugs, like Sildenafil, which enhances HCN activation, temporarily restore cognitive function without affecting the amyloid-beta load [28]. This suggests that alternate bio-chemical pathologies, apart from beta-amyloid plaques, can explain the early impairment of cognition. Alpha-rhythm disruption is reported in early stage Alzheimer’s [13]), along with a loss in spatial cognitive abiltites. The hippocampal theta-rhythm is known to play a crucial role in learning and spatial navigation [29] [30] [31]. The theta rhythm has also been shown to have critical dependence on HCN channels. [32] These investigations, taken together, point towards the need to investigate the the possibility of an HCN channel pathology in early stage Alzheimer’s.

We explore the space of all possible causal relationships between amyloid-beta(*Aβ*), HCN channel expression(H) and alpha-rhythms(*α*) to explore the various correlations that have been reported in literature. We have established that lowering the HCN channel expression reduces the alpha-rhythm frequency and coherence 3. The first box in 7 lists all the possibilities, where it is not the case that the alpha-rhythm(*α*) is effected by I_H_ current channels(H). Given the insights from our model (monotonic relationship between HCN expression and alpha peak frequency 3), these relationships are eliminated. In studies where the HCN channels were knocked out, Saito et al. report increased amyloid-beta beta aggregation when compared to wild type (WT) neurons [17]. They also show that using an HCN channel blocker(ZD7288) in WT also leads to similar levels of amyloid-beta accumulation as the KO neurons. Postmortem studies of AD patient brains have lower levels of HCN channels, when compared to non-AD subjects [17]. Using these observations, the relationships described in the second box, 7, can be eliminated. Revelations from our model combined with experimental observations shrinks the list of relationships to merely three possible causal links and is illustrated in the third box in 7. HCN channel expression effects both the alpha rhythm and Amyloid-*β* directly. However it is not not clear if there is a directional direct link between amyloid-*β* and the alpha rhythm.

**Fig 7.**
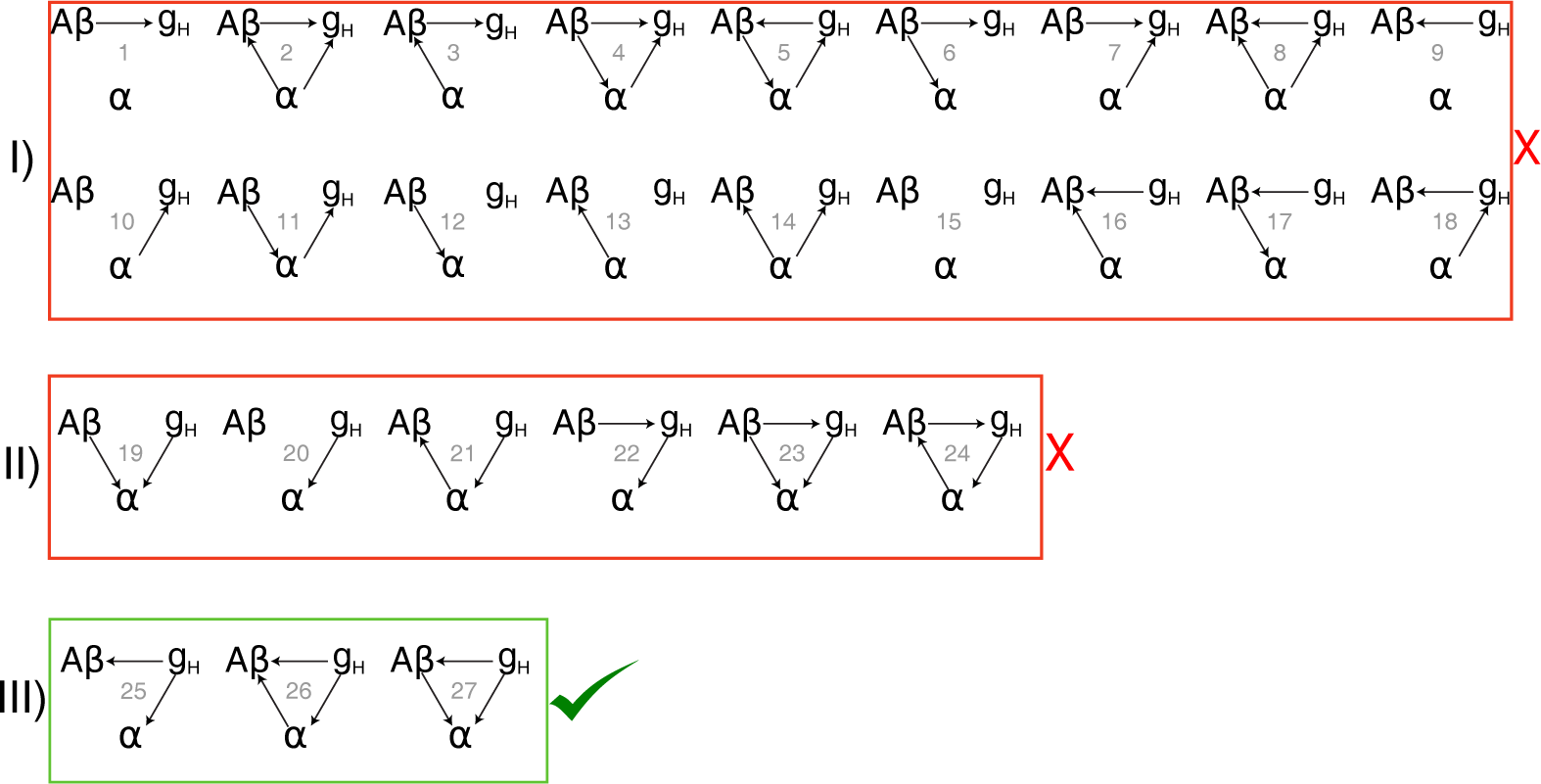
Potential causality between amyloid-beta plaques (*Aβ*), HCN channels(g_H_) and the alpha-rhythm(*α*). I)HCN expression directly effects the alpha-rhythm: elimination of the possibilities where this is not the case. II): Appearance of beta-amyloid plaques and lowered expression of HCN channels are strongly correlated and therefore not independent pathologies. It has been established that HCN channels activity effects amyloid-beta: elimination of box II possibilities [17]. III) Three remaining possibilities in the causal relationships

#### Mechanisms for loss in coherence

We characterize two distinct mechanisms that lead to loss in coherence and lower power in alpha as observed in AD. First we describe coherence loss with reduced g_H_ (3 g_H_ < 0.26 mS/cm^2^).

Since rise in I_H_ dictates the inter-burst interval, lower g_H_ increases the ISI and extends the the time spent by the membrane voltage close to the threshold of high threshold calcium channels. This increases the probability of background noise to cross the threshold causing a noisier alpha. Second, loss in coherence with increased acetylcholine is due to increased cell excitability. We demonstrate how excessive cholinergic modulation in the thalamo-cortical network leads to a sudden loss of coherence 4A(Green *η_ach_* > 15%). Under these circumstances a random background signal has a lower barrier to cross the threshold. This way the exact time of initiating the burst underlying the alpha rhythm becomes unreliable. The same mechanism also underlies the loss in coherence with increasing g_H_(3 g_H_ > 0.43mS/cm^2^). Under healthy physiological conditions, the HCN expression lies within a sweet spot, in a regime where it does not spend too much time near the calcium current threshold, and at the same time has a substantial barrier to cross to reach the threshold. This allows for a robust 10 Hz burst to be precisely orchestrated, that is predominantly unaffected by noise.

#### Alpha-rhythm relation to overall firing rates and extracellular GABA

A recent AD study reports abnormally high levels of the neurotransmitter GABA in the extracellular space [22] implicating enhanced GABAergic drive in the pathology. In support of this, temporary rescue of cognition in mice by reducing the inhibitory effect of GABA is reported. [22] Our model illustrates clearly that higher GABA levels can be a direct down stream effect of reduced HTC firing frequency and lower alpha rhythm (See figure 5). Our model predicts that higher GABA levels, a downstream effect of a slower alpha-frequency in AD may further result in a runaway affect of slowing down activity and exacerbate pathology 5.

The alpha-rhythm is often associated with a suppression of overall neuronal firing rates. [21] A lower alpha then would imply release from this suppression and increase in the overall firing rates. Our model provides an insight into the paradoxical effect of release from inhibition and increase in GABA. Reduction in HCN expression, that mimics AD pathology, corals other neurons in the network to enhanced activity and changes the global firing rates. 5 Around half the neurons in the network are GABA releasing RE cells. They show an increase in their activity, along with the glutamatergic TC neurons. This shows how reduced alpha-frequency can lead to both increased GABA levels and increased neural firing rates.

## Conclusion

Using an alpha rhythm generating network model of the thalamus and its disruption in AD, we have systematically elucidated the causal links between various known pathologies associated with Alzheimer’s Disease. We hypothesize that HCN pathology precedes alpha rhythm disruption and may underly early cognitive deficiency in the disease. Our results illustrate limitations of therapeutic intervention of enhancing acetylcholine and downstream effects of enhanced GABA activity. Mimicking increased calcium flux as seen in AD results in global changes in network firing rate and loss of coherence. When the HCN pathology is simulated, the AD network is overtly sensitive to changes in calcium signaling. Changes in brain rhythm is an early pathological signature in AD, this paradigm can contribute to our understanding of a nodal cause of the disease.

## Acknowledgments

SN funded by Wellcome DBT India Alliance intermediate fellowship and Indian Institute of Science Education and Research - Pune. RS is funded by Department of Science and Technology INSPIRE fellowship, Indian Institute of Science Education and Research - Pune and Wellcome DBT India Alliance.

## Supporting Information

**Fig 8.**
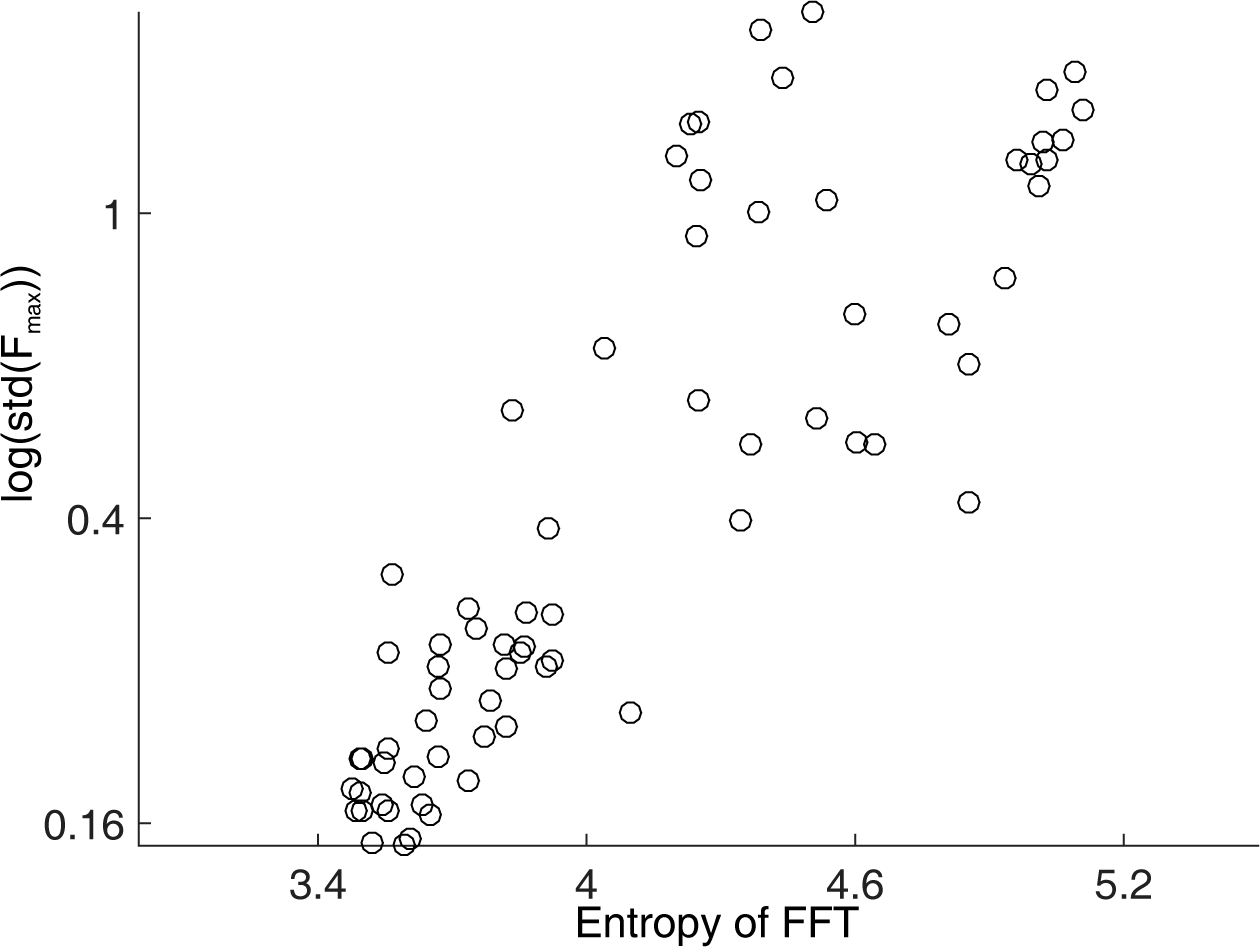
Standard deviation of the peak frequency increases with increase in entropy of FFT. Illustration of the relationship between entropy of FFT with the standard deviation of the peak frequency. The log of the standard deviation is plotted against the entropy.

## References

1. Rihs TA, Michel CM, Thut G. Mechanisms of selective inhibition in visual spatial attention are indexed by *α*-band EEG synchronization. European Journal of Neuroscience. 2007;25(2):603–610.

2. Buzsaki G. Rhythms of the Brain. Oxford University Press; 2006. Available from: https://books.google.co.in/books?id=ldz58irprjYC.

3. Fisahn A, Pike FG, Buhl EH, Paulsen O. Cholinergic induction of network oscillations at 40 Hz in the hippocampus in vitro. Nature. 1998;394(6689):186–189. Available from: http://www.nature.com/doifinder/10.1038/28179.

4. Lörincz ML, Crunelli V, Hughes SW. Cellular dynamics of cholinergically induced alpha (8-13 Hz) rhythms in sensory thalamic nuclei in vitro. The Journal of neuroscience: the official journal of the Society for Neuroscience. 2008;28(3):660–671.

5. Bollimunta A, Chen Y, Schroeder CE, Ding M. Neuronal Mechanisms of Cortical Alpha Oscillations in Awake-behaving Macaques. The journal of Neuroscience. 2009;28(40):9976–9988.

6. Rapid distributed fronto- parieto-occipital processing stages during working memory in humans. Cereb Cortex. 2002;12:710–728.

7. Magnetoencephalographic 10-Hz rhythm from the human auditory cortex. Neuroscience Letters. 1991;129:303–305.

8. Spatiotemporal characteristics of sensorimotor neuro-magnetic rhythms related to thumb movement. Neuroscience. 1994;60:537–550.

9. Haegens S, Nácher V, Luna R, Romo R, Jensen O. *α*-Oscillations in the monkey sensorimotor network influence discrimination performance by rhythmical inhibition of neuronal spiking. Proceedings of the National Academy of Sciences of the United States of America. 2011;108(48):19377–82. Available from: http://www.pnas.org/content/108/48/19377.abstract.

10. Vijayan S, Kopell NJ. Thalamic model of awake alpha oscillations and implications for stimulus processing. Proceedings of the National Academy of Sciences. 2012;2012:1–6.

11. Krishnan GP, Chauvette S, Shamie I, Soltani S, Timofeev I, Cash SS, et al. Cellular and neurochemical basis of sleep stages in the thalamocortical network. eLife. 2016;5(November 2016):1–29.

12. Friston KJ, Bastos AM, Pinotsis D, Litvak V. LFP and oscillations-what do they tell us? Current Opinion in Neurobiology. 2015;31:1–6.

13. Vitiello MV. Dominant occipital (alpha) rhythm frequency in early stage Alzheimer’s disease and depression. Sciences-New York. 1989;98195:427–432.

14. Crunelli V, David F, Lo˝rincz ML, Hughes SW. The thalamocortical network as a single slow wave-generating unit. Current Opinion in Neurobiology. 2015;31:72–80. Available from: http://linkinghub.elsevier.com/retrieve/pii/S0959438814001792.

15. Kunz L, Schröder TN, Lee H, Montag C, Lachmann B, Sariyska R, et al. No Title. Science. 2015;350(6529):430–433. Available from: http://www.sciencemag.org/content/350/6259/430.full.html.

16. Giocomo LM, Hussaini SA, Zheng F, Kandel ER, Moser MB, Moser EI. Grid cells use HCN1 channels for spatial scaling. Cell. 2011;147(5):1159–1170. Available from: http://dx.doi.org/10.1016/j.cell.2011.08.051.

17. Saito Y, Inoue T, Zhu G, Kimura N, Okada M, Nishimura M, et al. Hyperpolarization-activated cyclic nucleotide gated channels: a potential molecular link between epileptic seizures and A*β* generation in Alzheimer’s disease. Molecular neurodegeneration. 2012;7(1):50. Available from: http://www.molecularneurodegeneration.com/content/7/1/50.

18. Zou X, Coyle D, Wong-Lin K, Maguire L. Beta-amyloid induced changes in A-type K + current can alter hippocampo-septal network dynamics. Journal of Computational Neuroscience. 2012;32(3):465–477.

19. McCormick DA, Prince DA. Acetylcholine induces burst firing in thalamic reticular neurones by activating a potassium conductance. Nature. 1986;319(6052):402–405.

20. Vitiello G. The use of many-body physics and thermodynamics to describe the dynamics of rhythmic generators in sensory cortices engaged in memory and learning. Current Opinion in Neurobiology. 2015;31:7–12. Available from: http://dx.doi.org/10.1016/j.conb.2014.07.017.

21. Klimesch W. Alpha-band oscillations, attention, and controlled access to stored information. Trends in Cognitive Sciences. 2012;16(12):606–617. Available from: http://dx.doi.org/10.1016/j.tics.2012.10.007.

22. Wu X, Foster DJ. Hippocampal replay captures the unique topological structure of a novel environment. The Journal of Neuroscience. 2014;34(19):6459–69. Available from: http://www.ncbi.nlm.nih.gov/entrez/query.fcgi?cmd=Retrieve{&}db=PubMed{&}dopt=Citation{&}list{_}uids=24806672$\delimiter026E30F$nhttp://www.jneurosci.org/content/34/19/6459.abstract.html?etoc.

23. Berridge MJ. Calcium regulation of neural rhythms, memory and Alzheimer’s disease. The Journal of physiology. 2014;592(Pt 2):281–93. Available from: http://www.pubmedcentral.nih.gov/articlerender.fcgi?artid=3922493{&}tool=pmcentrez{&}rendertype=abstract.

24. Binder LI, Guillozet-Bongaarts AL, Garcia-Sierra F, Berry RW. Tau, tangles, and Alzheimer’s disease. Biochimica et Biophysica Acta - Molecular Basis of Disease. 2005;1739(2):216–223.

25. Selkoe DJ. Alzheimer’s Disease–Genotypes, Phenotype, and Treatments. Science. 1997;275(5300):630–631. Available from: http://science.sciencemag.org/content/275/5300/630.

26. Kumar A, Singh A, Ekavali. A review on Alzheimer’s disease pathophysiology and its management: An update. Pharmacological Reports. 2015;67(2):195–203. Available from: http://dx.doi.org/10.1016/j.pharep.2014.09.004.

27. Hughes SW, Crunelli V. Thalamic mechanisms of EEG alpha rhythms and their pathological implications. The Neuroscientist: a review journal bringing neurobiology, neurology and psychiatry. 2005;11(4):357–372.

28. Cuadrado-Tejedor M, Hervias I, Ricobaraza A, Puerta E, P´erez-Roldán JM, Garćıa-Barroso C, et al. Sildenafil restores cognitive function without affecting *β*-amyloid burden in a mouse model of Alzheimer’s disease. British Journal of Pharmacology. 2011;164(8):2029–2041.

29. Kleshchevnikov aM. Synaptic plasticity in the hippocampus during afferent activation reproducing the pattern of the theta rhythm (theta plasticity). Neuroscience and behavioral physiology. 1999;29(2):185–96. Available from: http://www.ncbi.nlm.nih.gov/pubmed/10432508.

30. Larson J, Wong D, Lynch G. Patterned stimulation at the theta frequency is optimal for the induction of hippocampal long-term potentiation. Brain Research. 1986;368(2):347–350. Available from: http://www.sciencedirect.com/science/article/pii/0006899386905792.

31. Eva Pastalkova, Vladimir Itskov, Asohan Amarasingham, and Gyorgy Buzsaki. Internally Generated Cell Assembly Sequences in the Rat Hippocampus. Science. 2008;321(5894):1322–1327.

32. Varga V, Hangya B, Kránitz K, Ludányi A, Zemankovics R, Katona I, et al. The presence of pacemaker HCN channels identifies theta rhythmic GABAergic neurons in the medial septum. Journal of Physiology. 2008;586(16):3893–3915.

